# Deconvolving the contributions of cell-type heterogeneity on cortical gene expression

**DOI:** 10.1101/566307

**Authors:** Ellis Patrick, Mariko Taga, Ayla Ergun, Bernard Ng, William Casazza, Maria Cimpean, Christina Yung, Julie A Schneider, David A Bennett, Chris Gaiteri, Philip L De Jager, Elizabeth M Bradshaw, Sara Mostafavi

## Abstract

Complexity of cell-type composition has created much skepticism surrounding the interpretation of brain bulk-tissue transcriptomic studies. We generated paired tissue genome-wide gene expression data and immunohistochemistry data, enabling us to assess statistical methods for modeling and estimating cellular heterogeneity in the brain. We demonstrate that several algorithms that rely on single-cell and cell-sorted data to define cell marker gene sets yield accurate *relative* and *absolute* estimates of constituent cell-type proportions.

## Introduction

The observed gene expression levels in tissues with high cellular heterogeneity are influenced by the proliferation or death of specific cell-types and also by molecular processes within cell-types. In the context of disease studies, this ambiguity in the origin of gene expression changes can generate spurious disease associations or reduce statistical power to detect true associations^1^. Accurately separating out the contributions of cell-type composition on gene expression, through a mathematical process known as deconvolution, should result in more accurate molecular measures of disease in heterogeneous tissue. This potential has been experimentally validated in specific settings, for instance on immune cell subsets^2^. Such approaches have been described for DNA methylation data in the brain to predict proportions of glial vs neuronal populations^3^.

Recent single-cell RNA-seq^4,5^ and cell-sorted datasets^6^ from human brain tissue can enhance the effectiveness of deconvolution methods through more accurate estimation of cell-type marker genes. Deconvolution algorithms are being adapted for application to gene expression in the brain using these cell markers to infer and adjust for glial cell subsets with higher granularity^7–9^. However, because of lack of availability of high-resolution benchmark datasets across multiple individuals, their accuracy and resolution is not well understood. Therefore, using a large cohort we have constructed a gold-standard brain dataset that can be used to contrast deconvolution method performance to estimate cell-type proportions and identify regulation within specific cell-types.

To establish a gold standard for cell-type proportions in heterogamous tissue, we used immunohistochemistry (IHC) to experimentally measure the proportion of neurons, astrocytes, microglia, oligodendrocytes and endothelial cells from dorsolateral prefrontal cortex (DLPFC) tissue of 70 older individuals. These samples are a subset of the larger ROSMAP cohort with bulk RNAseq (n=508) from same region^10^; donors showed a range of cognitive function, from healthy to Alzheimer’s dementia, which likely enhances the heterogeneity of cell-type proportions.

To generate IHC-based cell-type proportions, antibodies were chosen to identify neurons (NeuN), astrocytes (*GFAP*), microglia (*IBA1*), oligodendrocytes (*OLIG2*) and endothelial cells (*PECAM*). Automated image analysis was used to identify cells by DAPI staining and the cells that were positive for a particular cell-type marker (**Figure 1A**). Testing the quality of the IHC data, first, we observed that the proportion of the five major cell populations per subject approximately sums to one, despite separate staining for each cell-type marker (**Figure 1B**). In addition to indicating the accuracy of the counts, this observation also implies that the five cell-types measured make up the bulk of the DLPFC, and no major population is unmeasured. Second, the IHC estimates correlate with expression levels of gene modules that are enriched for cell-type specific markers that were previously defined from this data^11^ (**Figure S1**). In total, IHC-based estimates of cell-type proportions explained ~8% of the variation in gene expression levels, indicating the data is a relevant testbed for deconvolution, as many genes correlate with the heterogeneity in cellular proportions (**Figure 1C**).

**Figure 1.**
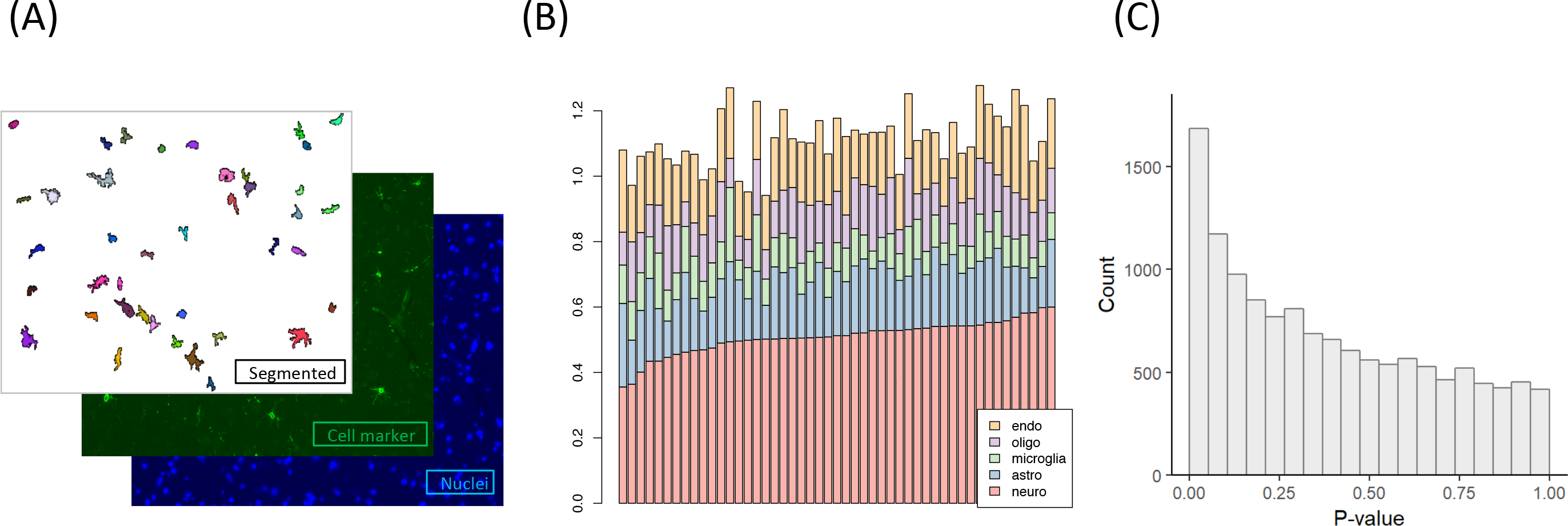
Estimation of cell type proportions by IHC. (A) Figure depicts example images used to quantify cell type proportions. (B) Each bar represents an individual, y-axis shows the estimated proportion of each of the five cell types. (C) P-value distribution, showing the p-values for the correlation between gene expression levels (all expressed genes) and IHC-based cell type proportions estimates across 70 individuals with paired data.

Using the IHC data as a standard, we compared the accuracy of popular deconvolution methods. Methods fell into two classes: 1) “supervised” reference-based methods, which included non-negative least squares (NNLS)^12^, CiberSort^13^ and dtangle ^7^ and 2) “semi-supervised” reference-based, exemplified by DSA^14^. Both classes rely on pre-defined marker genes (also referred to as signature gene lists) for each cell-type; the distinction is that supervised approaches also require cell-specific *expression profiles* (derived from cell-specific gene expression datasets) for the marker genes.

In conjunction with the methods comparison, we used three typical sources for cell-type marker genes: (1) human single-cell RNA-seq data^15^, (2) human cell-sorted RNA-Seq data^16^, and (3) a curated collection of cell-sorted microarray data and In-Situ Hybridization from mouse (Neuroexpresso)^17^. For each marker gene data source, differential gene expression analysis identified sets of marker genes that are preferentially expressed in each of the five cell-types. Results for a given method were consistent across different sources of marker genes, with greatest variability in the estimates for endothelia and microglia (**Figure S2**). For simplicity we focus on results from single-cell RNA-Seq based markers (others shown in supplement).

We assessed the concordance between IHC estimates and deconvolution algorithms in two ways: 1) based on the correlation between the inferred and measured *relative* proportions for each cell-type across individuals and 2) based on the population-level *absolute* proportion across cell-types. Four trends emerge from these analyses. First, correlations between IHC and deconvolution estimates were typically significant, with moderate effect sizes, but variable results for endothelial cell proportions (**Figure 2A, S3A**). Secondly, we observe the importance of robust multi-gene markers for accurate deconvolution. Specifically, the endothelial results point to noise in the available signature gene sets, as single-cell-based defined marker genes for endothelial cells performed worse than those defined based on cell-sorted data and the semi-supervised approach (**Figure 2A, S1, S3**). At the same time, we find potentially weaknesses in ‘single marker’ approaches, as *ENO2* typically used for approximating the proportion of neurons is not predictive of the overall proportions of neurons, as compared to estimates provided by deconvolution algorithms. Third, the various algorithmic approaches yield highly correlated estimates as assessed more robustly across a larger set of 508 ROSMAP samples (**Figure S4**). However, Cibersort and NNLS were “outliers” in this respect for estimation of microglia cells, which may stem from their difficulty in estimating such low abundant cell-types (**Figure 2, S3**). Fourth, of practical importance, we observed that IHC proportions across cell-types were highly concordant with *absolute* proportions estimated by the deconvolution algorithms (**Figure 2B, S3B**), with NNLS generally providing the worst performance. The across cell-type concordance implies that the estimated proportions are not confounded by the variability in the total amount of RNA across different cell-types, as one may suspect. We also assessed the robustness of these results with respect to variability in marker gene set size, and found the results to be robust for a wide range (**Figure S5**).

**Figure 2.**
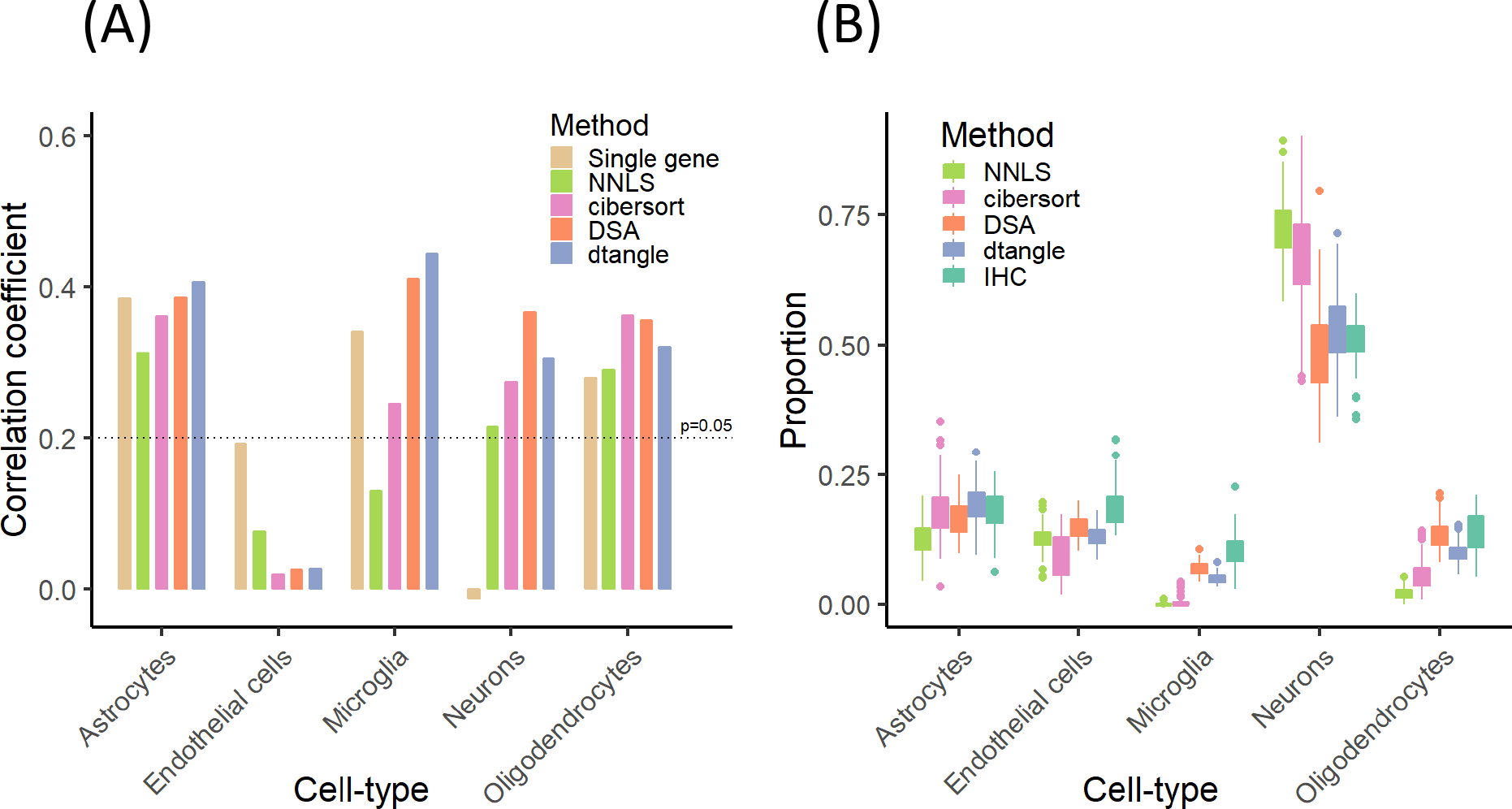
Comparison of deconvolution algorithms. (A) Figure shows the Pearson correlation coefficient between IHC-based cell type estimate and four deconvolution algorithms, in addition to the “single marker” based approach. For the single marker based approach, we used the expression of the widely used marker genes: ENO2 for neurons, GFAP for astrocytes, CD68 for microglia, CD34 for endothelial, OLIG2 for oligodendrocytes. (B) Estimates of *absolute* proportions of each cell types according to the four algorithms tested, and IHC (experimentally measured in this study). Box plots depict the range of proportions across 70 individuals. For both (A) and (B),

Additionally, we applied the same approach to predict cell-type proportions across 9 brain regions based on GTEx data^18^, with prediction that cell-type proportions vary strongly across these nine regions (**Figure S6**), with adjacent regions tending to yield similar proportions, which indicates the stability of the methods. Although not much is conclusively known about the variation in cell type proportions across human brain regions^19^, encouragingly, when data were available, these predictions matched what was expected based on cell counts using single-cell RNA-seq data^4^ (**Figure S7**).

To demonstrate the utility of cell-type deconvolution in the brain, we used the predicted cell-type proportions to perform cell-type-specific eQTL analysis^20^. First, we hypothesized that deconvolution algorithms that utilize groups of marker genes should yield more accurate prediction of cell-type proportions, and hence increase the statistical power for cell-type specific eQTL analysis compared to single marker type approaches. Indeed we confirmed a significant gain in sensitivity in detecting cell-type specific eQTLs when we used deconvolution algorithms as opposed to single markers (**Figure 3A**). For instance, single marker based proxies of cell-types produced 7 cell-type specific eQTLs, while DSA produced 232. As one example, we found SNPs near STMN4 were significantly associated with its expression but the correlation was dependent on the proportion of oligodendrocytes (**Figure 3B**). Fittingly, STMN4 is highly expressed in oligodendrocytes.

**Figure 3.**
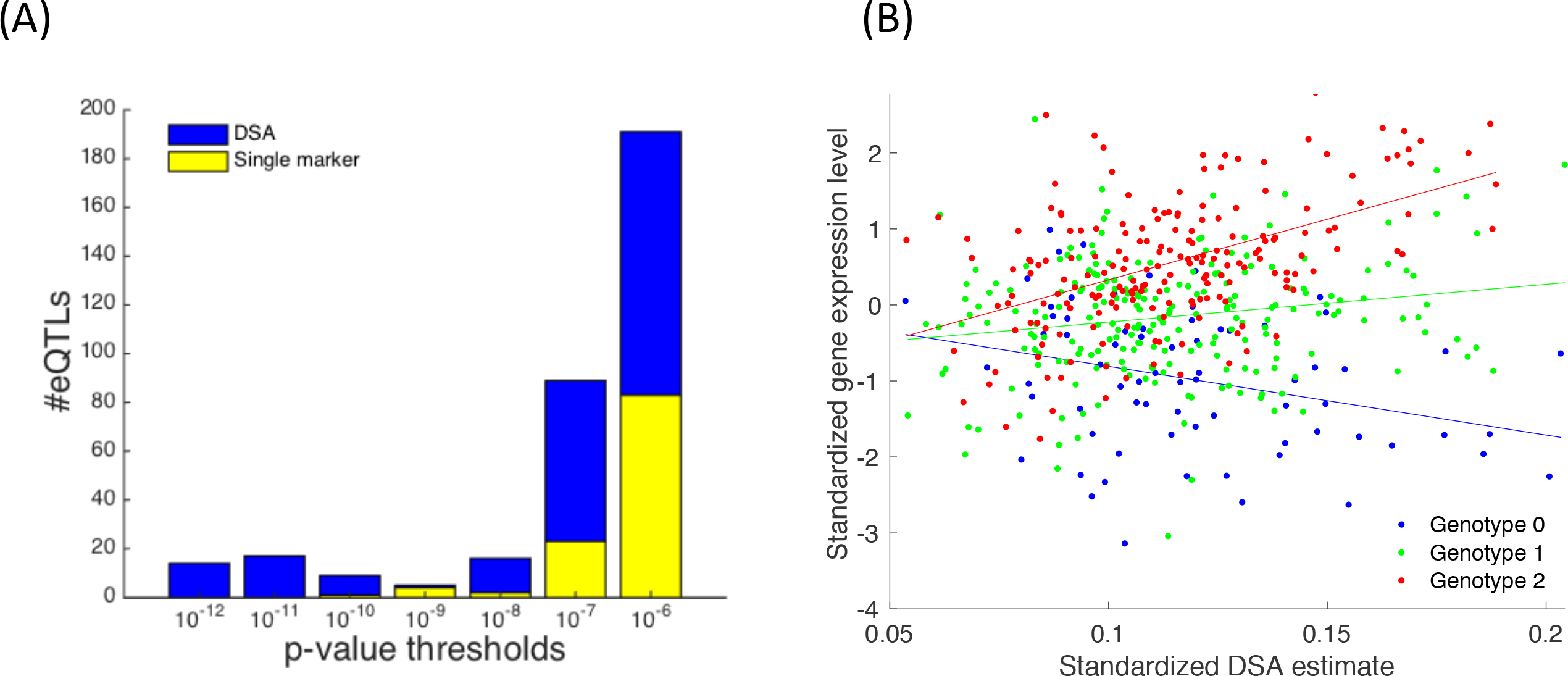
Discovery of cell-type specific eQTLs. (A) Figure shows the number of associations for several p-value thresholds. Number of associations found based on the DSA estimates are shown in blue, and those based on single cell marker genes are shown in yellow. (B) An example of cell-type specific eQTL for oligodendrocytes. Figure shows the relationship between proportion of oligodendrocytes (as predicted by DSA) and expression levels of the STMN4 gene. The different colors depict the different genotype groups (0,1,2) for the associated SNP rs10481349.

In summary, we generated IHC data and used image analysis to quantify cell-type proportions in the brain. This provided an independent dataset for validation of cell-type deconvolution algorithms for bulk brain transcriptomic data. Our analysis concludes that several deconvolution algorithms yield predictions that are significantly correlated with quantifiable cell-type proportions, and with each other.

## Supplementary Methods

### IHC image acquisition

Six μm sections of formalin-fixed paraffin embedded tissue have been stained for NeuN (Millipore), GFAP (Dako), Iba1 (Wako), Olig2 (Sigma) and PECAM-1 (Novus biologicals) using antigen retrieval Buffer (Citrate Buffer pH 6.0) for each marker. Sections have been blocked with blocking medium containing 3% BSA and incubated with primary antibodies for overnight at 4oC. Sections have been washed three times with PBS before incubation with Fluorophore-conjugated secondary antibody (Thermofisher) for one hour and coverslipped with anti-fading reagent containing Dapi (P36931, Life technology). Using fluorescence upright microscope (Zeiss Axio), 30 images have been captured in grey matter for each section at magnification x20 with a set exposure time in a systematic zigzag pattern to ensure that all layers of the cortex have been included in quantification.

### IHC image analysis

EBImage^21^ was used for all image analysis including background correction, thresholding and segmentation. Automated image analysis was used to identify cell nuclei by DAPI staining and the cells that were positive for a particular cell-type marker. For each particpant, proportions were estimated as the average proportion of cell marker positive nuclei across the replicate images. R scripts with the parameters used for estimating the proportions are located on https://github.com/ellispatrick/CortexCellDeconv as well as the corresponding IHC images.

### Defining cell-type markers

Three datasets were used to define marker genes and cell-type reference profiles. Cell-specific reference profiles were collected from single-cell RNA sequencing data (Darmanis)^15^ and RNA sequencing profiles of purified populations of cells (Zhang)^16^ and a set of curated markers from Neuroexpresso^17^. For Darmanis and Zhang, samples were TMM normalized and then voom^22^ was used to define marker genes. The markers were selected as the 100 genes with largest fold-change after filtering for genes with false discovery rate less than 0.05. (Performance with respect to varying marker set size is shown in Supplementary Figure S5.)

### Description of the Deconvolution Algorithms

Four cell-type deconvolution algorithms were applied to the data; Cibersort ^13^, dtangle ^7^, DSA ^14^ and NNLS ^12^. These algorithms were applied to the 508 RNA-seq samples from ROSMAP cohort, processed as previously described^11^. Briefly RNA-seq data was adjusted for known technical and biological factors, including age, sex, PMI, PH, and batch. For each of the deconvolution algorithm tested, we used the package provided as part of the primary paper and glmnet ^23^ was used for NNLS. Cibersort, dtangle and NNLS each require both cell-type reference profiles and marker genes while DSA just requires marker genes. For assessing correlations between gene expression and IHC, speakeasy clustering^24^, an unsupervised approach, was also evaluated using a set of predefined gene coexpression modules^10^ as well as the individual marker genes used in the IHC. As *CD31* wasn’t expressed in the gene expression data, *CD34* was used as the gene marker for endothelial cells instead. See above for the details of the marker set selection approach and https://github.com/ellispatrick/CortexCellDeconv for R scripts.

### Cell-type specific eQTL analysis

We used the approach described by Westra and colleagues^20^ to identify cell-type specific eQTLs. This approach tests for the statistical significance of a linear interaction model as follows:

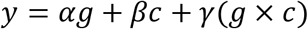

where *y* is a vector of gene expression levels, g is the genotype for the test SNP, *c* is the proportion of test cell-type, and *g x c* is the interaction term between genotype and the proportion of cell-type. The statistical significance of the interaction term, modeled by *γ*, implies the existing of a cell-type-by-genotype effect. As suggested by Westra and colleagues, to reduce the burden of multiple testing, only cis-SNPs previously found to be a *cis* xQTL (main effect), with a window of 1Mb around TSS, where tested. The cell-type estimates from the DSA algorithm where used. Global false discovery rate (FDR) threshold of 0.1 (correcting for all SNP-gene pairs and cell-types tested) was used to identify significant cell-type-by-genotype eQTLs.

