## Supplemental Figures for "Deconvolving the contributions of cell-type heterogeneity on cortical gene expression"

### Figure S1

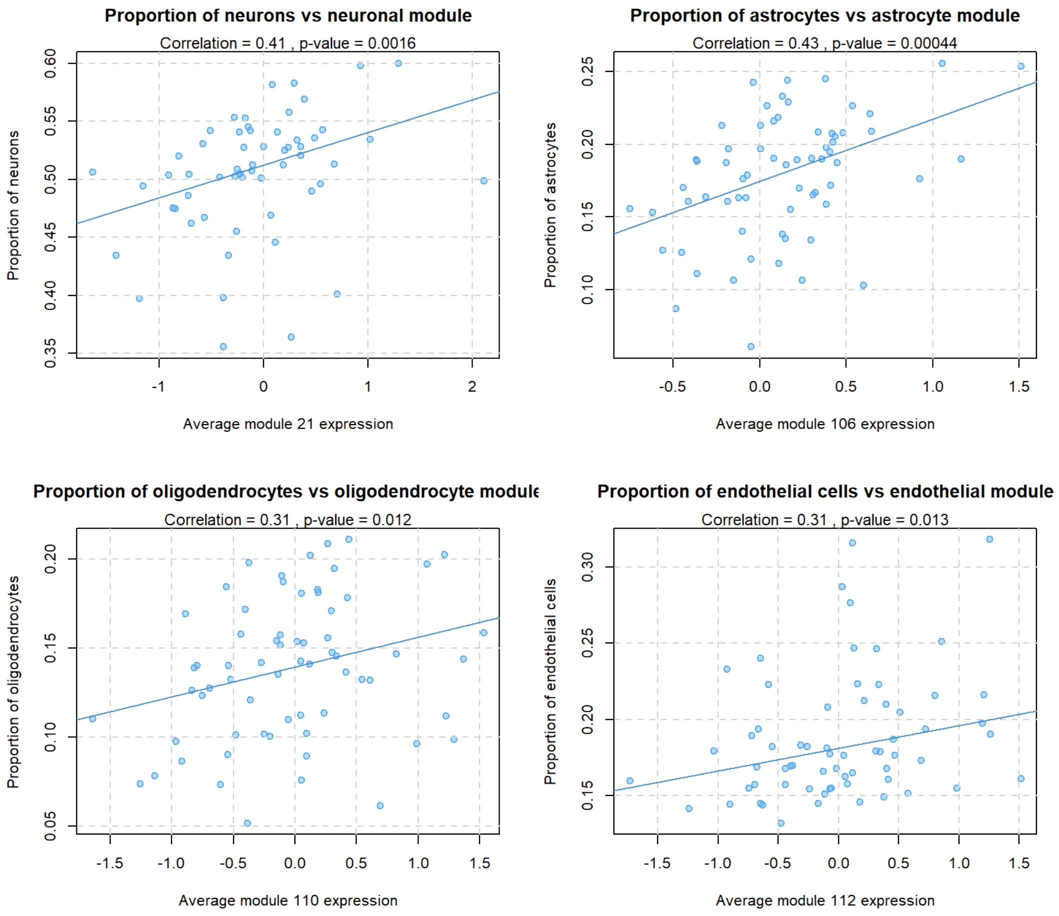


**Figure S1. Correlation between IHC estimates and expression level of gene modules.** Each dot depicts an individual. Our previous study defined a set of modules with gene members that were enriched for each of the five cell types examined (Mostafavi and Gaiteri et al., Nat Neur 2018): the average expression of each of these modules (across genes) represents a relative score for each individuals that can serve as a proxy for proportion of the corresponding cell type. The module average expression is shown on the x-axis and the IHC-based proportions are shown on the y-axis.

### Figure S2

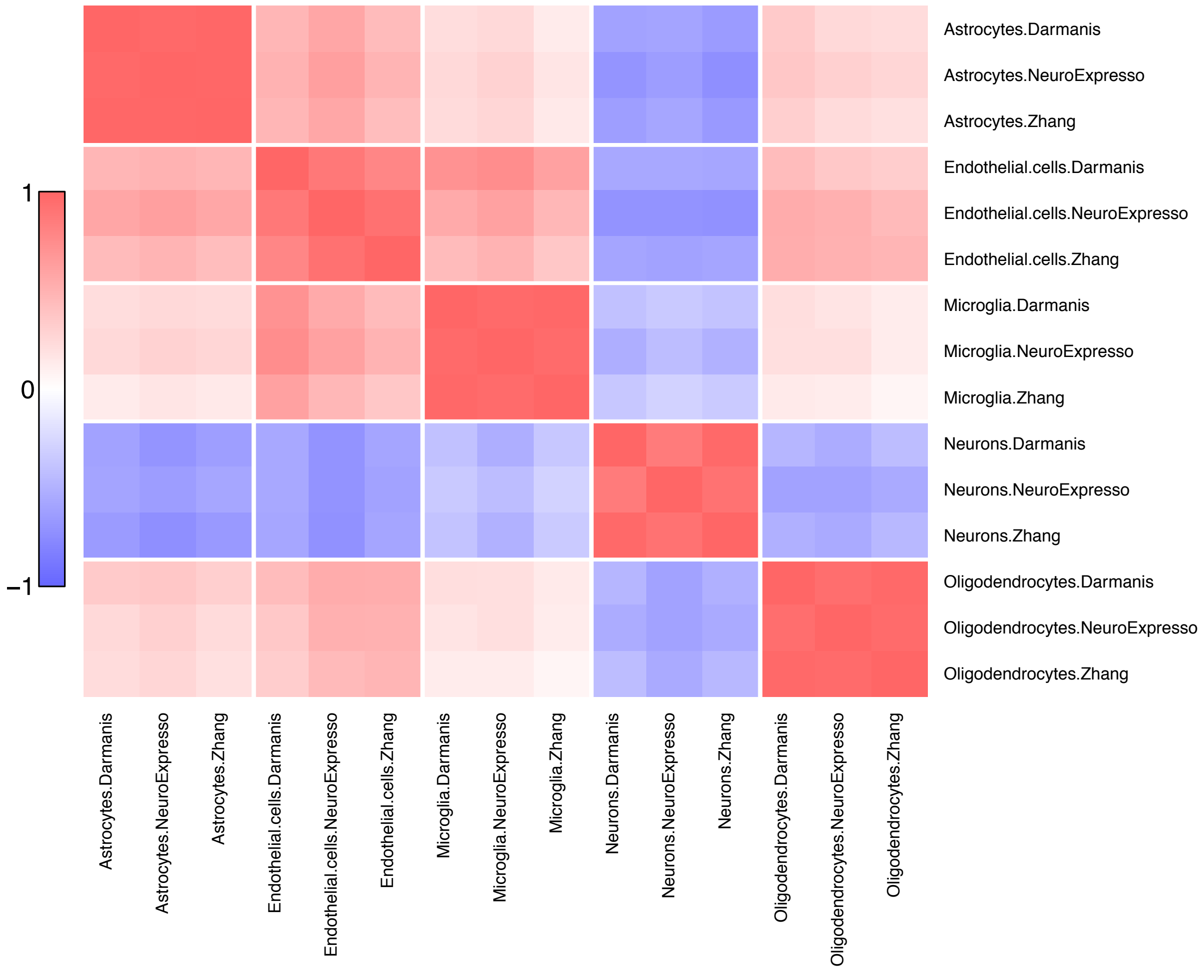


**Figure S2. Correlation between deconvolution algorithms with varying marker gene sets.** Three resources were used to define marker gene sets: 1) single cell RNA-seq data (Darmanis et al., Cell Reports 2017), 2) human cell-sorted data (Zhang et al., Neuron 2016), and 3) curated ISH data (NeuroExpresso; Mancarci et al., eNeuro 2017). Heatmap shows the Pearson correlation coefficient between the estimation of cell type proportions across 508 individuals by the DSA algorithm with varied source of marker gene sets.

Figure S3

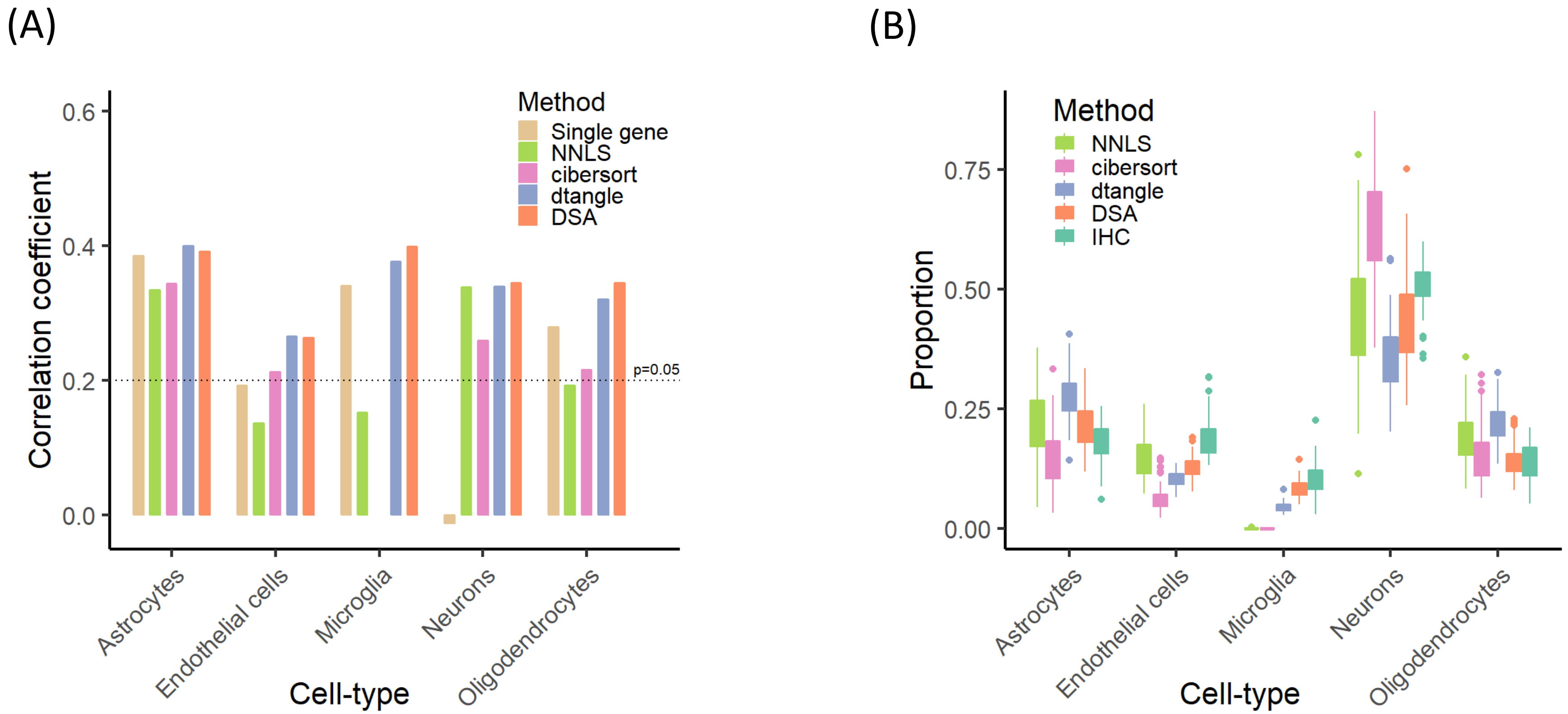


**Figure S3. Comparison of deconvolution algorithms based on marker genes defined on cell-sorted data.** (A) Correlation between prediction of cell type proportions using cell-sorted defined marker genes (Zhang et al, Journal of Neuroscience 2014) and IHC-based estimates. (B) Population-level range of predicted proportions of cell types using cell-sorted defined marker genes.

Figure S4

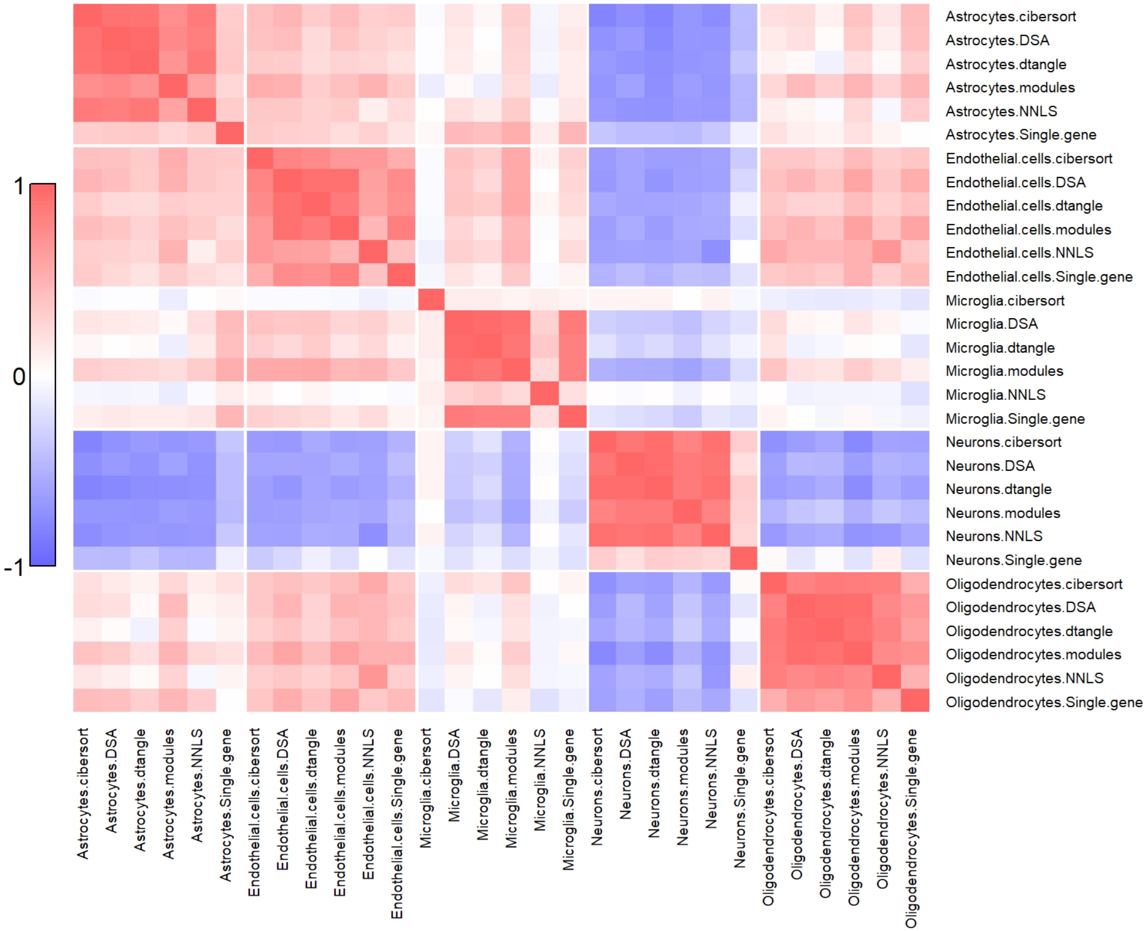


**Figure S4. Correlation of different deconvolution methods.** Plots show the pairwise correlation between pairs of deconvolution methods using the Zhang markers, assessed based on 508 samples.

Figure S5

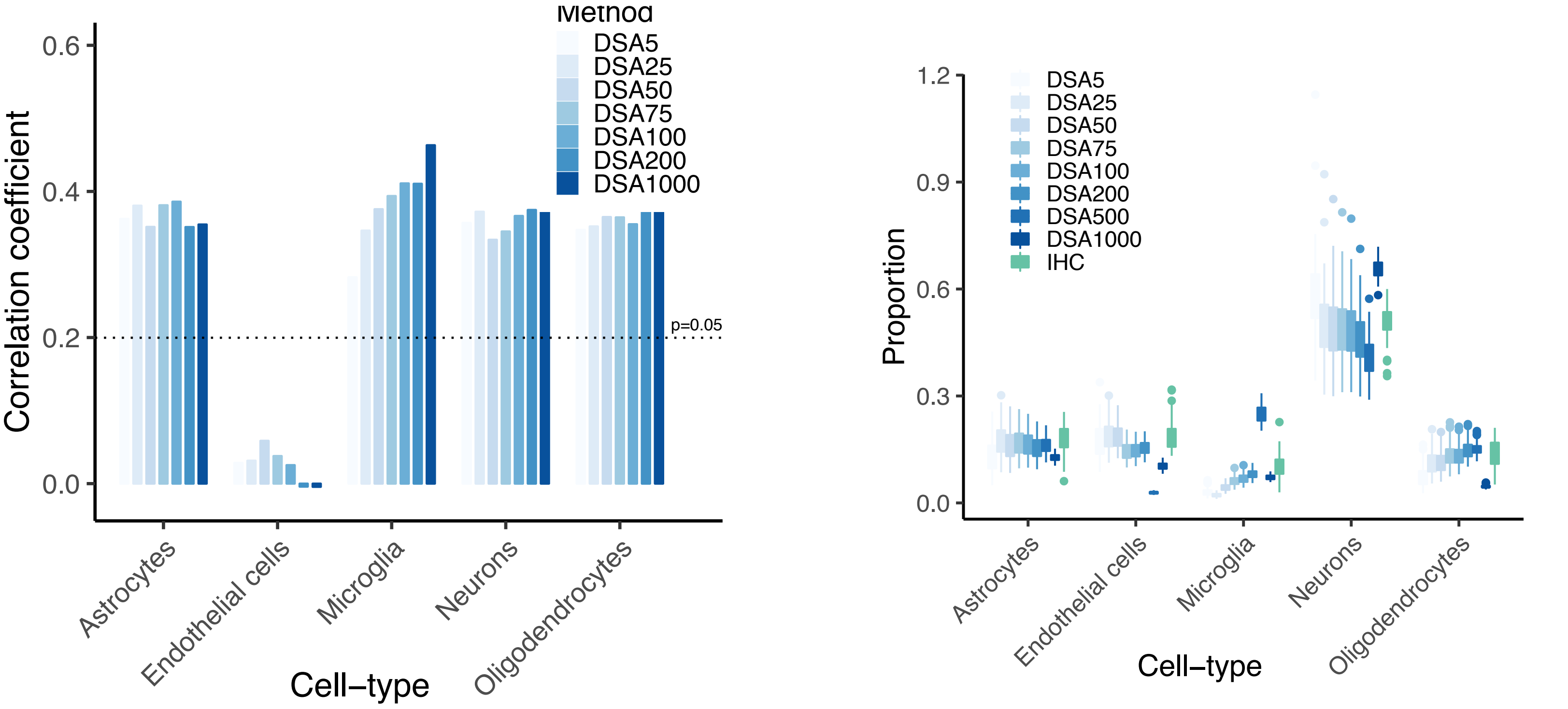


**Figure S5. Accuracy of predicted proportions with variable marker gene set size.** (A) Correlation between prediction of cell type proportions with variable sizes of marker gene sets. Differential expression analysis using single cell data (Darmanis et al., Cell Reports 2017) was used to define marker gene sets. (B) Population-level range of prediction of *absolute* proportions with variable size of marker gene sets.

### Figure S6

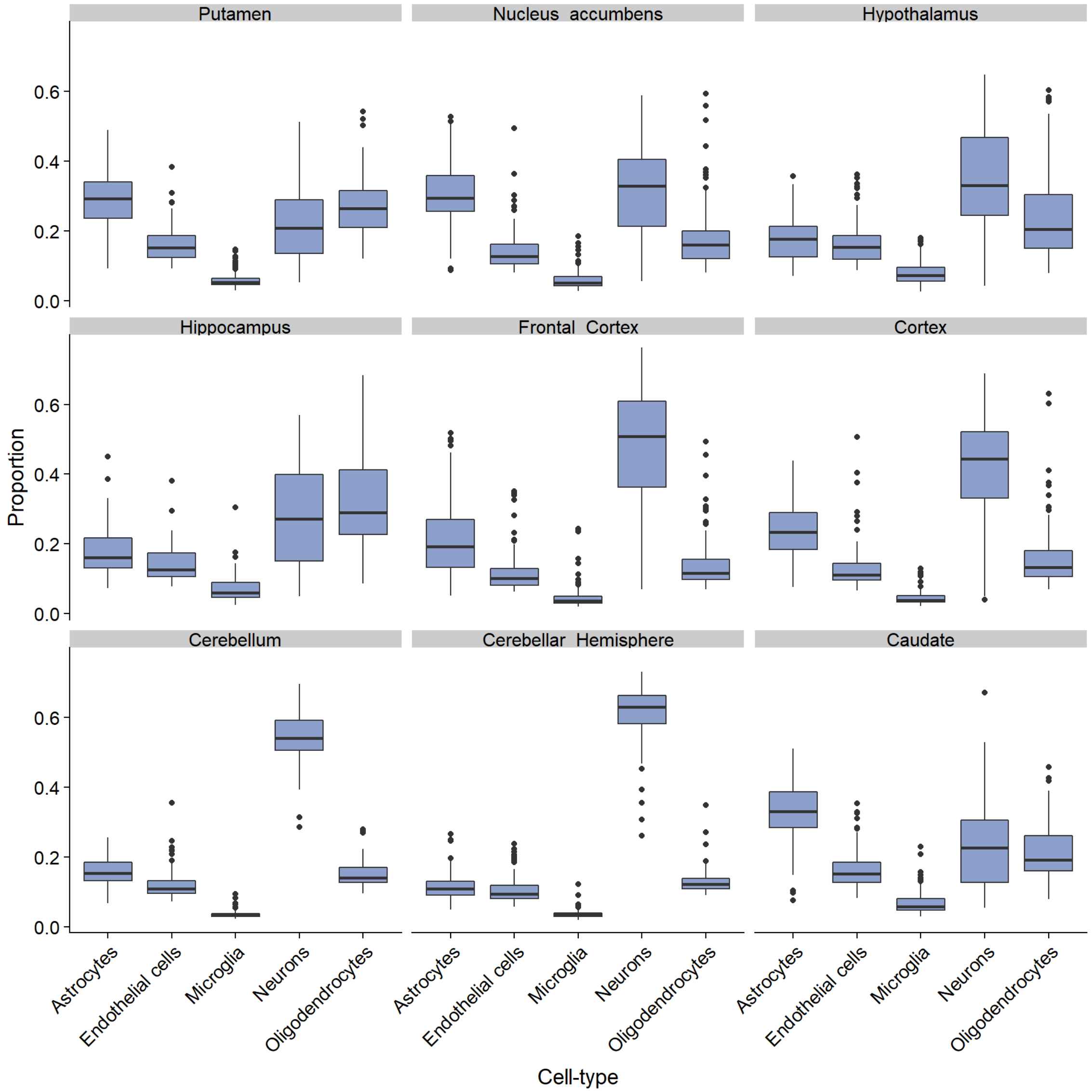


**Figure S6. Varied cell type proportions across human brain regions.** Cell type proportions were predicted, using the dtangle algorithm, across 9 brain regions using the GTEx bulk gene expression data.

### Figure S7

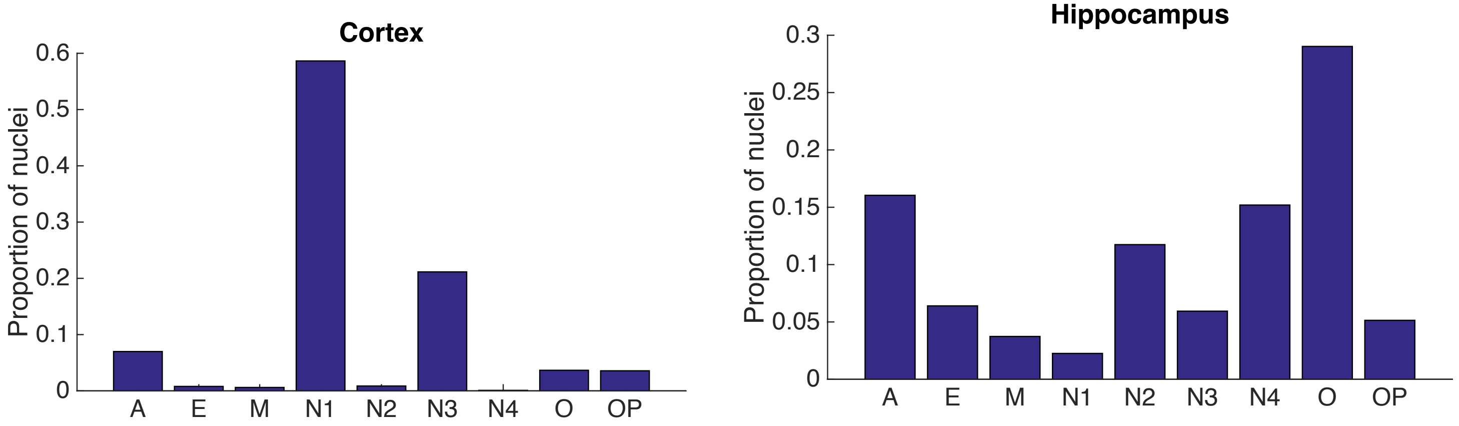


**Figure S7. Proportion of nuclei assigned to various cell types according to Dronc-Seq single-cell data from cortex and hippocampus.** Figure summarizes the proportion of nuclei assigned to various cell types (Habib et al., Nature Methods 2017). A: astrocytes; E: endothelial cells; M: microglia; N1,N2,N3,N4: different neuronal populations; O: oligodendrocyte; OP: oligodendrocyte progenitor cells.
